# Characterization of the Key Region and Phosphorylation Sites of EcaICE1 for its Molecular Interaction with EcaHOS1 Protein from *Eucalyptus camaldulensis*

**DOI:** 10.1101/861088

**Authors:** Ling Cheng, Weihua Zhang, Jianlin Hu, Ziyang Zhang, Yan Liu, Yuanzhen Lin

## Abstract

ICE1 (inducer of CBF expression 1), a MYC-like bHLH transcriptional activator, plays an important role in plant under cold stress via regulating transcriptional expression of downstream cold-responsive genes. Ubiquitination-proteasome pathway mediated by high expression of osmotically responsive gene1 (HOS1) can effectively induce the degradation of ICE1 and decrease the expression of expression of *CBFs* and their downstream genes under cold stress response in *Arabidopsis*, but the knowledge about ubiquitination regulation of ICE1 by HOS1 is still limited in woody plants. In this study, a complete open reading frame of *EcaICE1* gene and *EcaHOS1* was amplified from the tissue culture seedlings of *Eucalyptus camaldulensis*. Yeast two-hybrid (Y2H) and BiFC assay results showed that EcaICE1 can interact with EcaHOS1 protein in the nucleus, and further Y2H assay demonstrated that the 126-185 amino acid region at the N-terminus of EcaICE1 protein was indispensable for its interaction with EcaHOS1 protein. Moreover, we found that the amino acids at positions 143, 145, 158 and 184 within the key interaction region were the potential phosphorylation sites of EcaICE1 based on bioinformatics analysis, and that only the substitution of Serine (Ser) 158 by Alanine (Ala) blocked the protein-protein interactions between EcaICE1 and EcaHOS1 by Y2H and β-galactosidase assays using site-direct mutagenesis. In summary, we first reported that EcaICE1 could interact with EcaHOS1 protein in *Eucalyptus*, and identified Ser 158 of EcaICE1 as the key phosphorylation site for its interaction with EcaHOS1 protein.

## 1. Introduction

Cold stress is a major environmental factor that adversely affects plant growth and development, as well as the yield, product quality and geographic distribution of crops [1]. Plants must adjust various physiological and biochemical processes in response to cold stress by reprogramming gene expression [2]. Currently, the most well-understood cold signaling pathway is the ICE-CBF-COR transcriptional regulatory cascade [3,4]. CBF transcription factors (TFs) recognize the CRT/dehydration-responsive element (DRE) elements in the promoters of certain cold-responsive (*COR*) genes and regulate their expressions and functions [5]. Inducer of CBF expression 1 (ICE1), a MYC-like bHLH transcriptional activator, acts upstream as a positive regulator of CBFs by binding to MYC recognition elements in the *CBF* promoters in cold-responsive signaling [4,6,7]. In addition, *CBF* genes also appear to be negatively regulated by ICE1 via its interactions with MYB15 [8] and HOS1 [9], while positively by ICE1 via their interactions with SIZ1 [10] and OST1 [11]. Recently, Li et al. reported that MPK3/MPK6 could interact with and phosphorylate ICE1, reducing the stability of ICE1 as well as its transcriptional activity, thus negatively regulating CBF expression and freezing tolerance in *Arabidopsis* [12]. Therefore, ICE1 is the key regulators of ICE-CBF-COR transcriptional regulatory cascades in the cold-responsive signaling pathway. ICE-like genes have been isolated and characterized in some woody plants as *Populus suaveolens* [13], *Malus domestica* [14], *Vitis amurensis* [15], *E. camaldulensis* [16], *Pyrus ussuriensis* [17], *Poncirus trifoliata* [18] and *Hevea brasiliensis* [19]. However, the positive and negative regulation pathway of ICE1 in relation to cold stress response still remains poorly understood in woody plants.

*Eucalyptus* species are the important commercial woody plants because of rapid growth, broad adaptability, and the source of wood pulp. However, even widely distributed, its plantation is mainly restricted to tropical and subtropical regions because of freezing sensitivity, especially for the commercial *Eucalyptus* species. Therefore, it is necessary to discover the molecular regulation mechanism of cold response and carry out genetic improvement on freezing tolerance in *Eucalyptus*. Although *CBF* genes have been isolated and characterized from *Eucalyptus* [20,21,22,23,24,25], the knowledge about its upstream regulator ICE1 and its positive and negative regulation pathway needs to be investigated. Our precious studies have revealed that ectopic expression of *EcaICE1* from *E. camaldulensis* confers improved cold tolerance and the expression level of downstream genes in transgenic tobaccos [16]. Nevertheless, the factors controlling ICE1 protein stability associated with cold stress response in *Eucalyptus* are not clearly elucidated. In this study, we cloned the HOS1 gene *EcaHOS1* from the *E. camaldulensis* and analyzed the protein interactions between EcaICE1 and EcaHOS1 using Bimolecular fluorescence complementation (BiFC) and Yeast two-hybrid (Y2H) assays. Subsequently, the key phosphorylation sites of EcaICE1 for its interaction with EcaHOS1 protein were predicted by bioinformatics and characterized by site-direct mutagenesis using Y2H and assays.

## 2. Materials and methods

### 2.1. Plant materials and treatments

The tissue culture plantlets of *E. camaldulensis* cv. 103 were performed as described previously [16] and 30-day-old rooting plantlets were used in this study. Tobacco (*Nicotiana benthamiana*) plants were cultured in a growth chamber at 25 °C with 16/8 h light/dark photoperiod, and 5-week-old plants were used for further subcellular localization and BiFC analysis.

### 2.2. RNA isolation, gene cloning and sequence analysis

Total RNA was extracted as described previously [16], and treated with DNase I (Promega, Madison, WI, USA). 1 μg DNA-free total RNA was used the template for synthesizing the first strand cDNA (PrimeScript II 1st Strand cDNA Synthesis Kit; Takara, Dalian, China). The primers of *EcaICE1* and *EcaHOS1* (listed as table 1) were used for amplifying the aim genes with the first strand cDNA as the template. The aim genes were sequenced at Beijing genomics Institute (BGI, China). The coding sequence (CDS) of *EcaICE1* and *EcaHOS1* were predicted using FGENESH 2.6 software (http://linux1.softberry.com/berry.phtml?topic=fgenesh&group=programs&subgroup=gfind), and further confirmed by BLASTP program on the NCBI website (http://blast.ncbi.nlm.nih.gov/Blast.cgi). The protein secondary domains were predicted by Motif scan (https://myhits.isb-sib.ch/cgi-bin/motif_scan). Finally, sequence alignments with other plants were performed with CLUSTALX software.

**Table 1.**
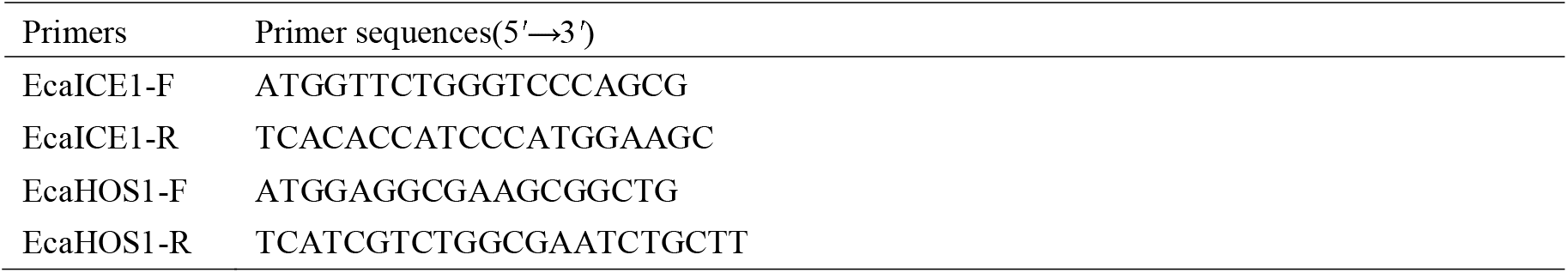
Primers used in the experiments.

### 2.3. Subcellular localization

The full-length CDS of *EcaICE1* was amplified by RT-PCR using primers (listed as Supplemental table S1) and fused into the 5’-terminus of the yellow fluorescent protein (YFP) of the pEarleyGate101 vector, driven by CaMV 35S promoter. The recombinant plasmid *35S::EcaICE1-YFP* was transferred into *Agrobacterium tumefaciens* strain GV3101 by heat shock method. The strain GV3101 harboring plasmid 35S::EcaICE1-YFP were cultured in the liquid yeast extract broth medium at 28□°C on a shaker at 220□rpm until the absorbance of the OD measurement at 660 nm reached 1.0~1.5, and then the culture were centrifuged at 8000 × g for 5□min. The thallus was mixed with the infecting solution containing 1% MES, 1% MgCl_2_ and 0.1% acetosyringone, and cultured for 3□h at 28°C and then injected into the abaxial surfaces of 5-week-old tobacco leaves via a syringe with an incubator for 48-72 h. The YFP fluorescence in the tobacco leaves was visualized using a confocal microscopy (Zeiss LSM 710, Carl, Germany) with excitation and emission at 513 and 527 nm, respectively.

### 2.4. Bimolecular fluorescence complementation (BiFC) assay

To perform BiFC assays, the whole CDS of *EcaICE1* and *EcaHOS1* (without their stop codons) were subcloned into pUC-pSPYNE or pUC-pSPYCE vectors as described previously [26] using primers (listed as Supplemental table S2). Expressions of each target gene alone were used as negative controls. The recombinant vectors were used for transient assays of tobacco leaves as described earlier. The transformed tobacco leaves were then kept in an incubator at 22 °C for 24-48 h. YFP signal was examined using a confocal microscope (Zeiss LSM 710, Carl, Germany).

### 2.5. Yeast two-hybrid (Y2H) assay

Yeast two-hybrid assays were carried out using the Matchmaker™ gold Yeast two-Hybrid Systems (Clontech, USA). Different truncated CDS of *EcaICE1* without transcriptional activation activity were subcloned into pGBKT7 (BD) to form the bait vector. The full-length CDS of *EcaHOS1* was amplified and inserted into pGADT7 (AD) to construct the prey vector (AD-*EcaHOS1*). The primers are listed in Supplemental table S3. The bait and prey vectors were co-transformed into yeast strain gold Y2H using lithium acetate (LiAc) method. Then yeast cells were plated on SD/-LW medium (minimal media double dropouts, SD basal medium without Leu and Trp) according to the manufacturer’s protocol (Clontech, USA) for 72 h. Transformed colonies were sprayed on SD/-LWHA medium (minimal media quadruple dropouts, SD medium with -Leu/-Trp/-Ade/-His) containing 125-μM Aureobasidin A (AbA), to test for possible protein-protein interactions, according to the yeast cell growth status. Each experiment replicated three technological repeats in separate experiments.

### 2.6. Phosphorylation sites prediction and site-direct mutagenesis

The phosphorylation sites within the key region of EcaICE1 for its interaction with EcaHOS1 were predicted using NetPhos 3.1 server (http://www.cbs.dtu.dk/services/NetPhos/), and further analyzed by ProtParam program on the expasy website (http://web.expasy.org/protparam). The potential phosphorylation sites were used for substitution into alanine by site-direct mutagenesis. Site-direct mutagenesis experiments were carried out with KOD-Plus Mutagenesis Kit (TOYOBOCO, China). The BD-EcaICE1_T3_ plasmid was used as the temple for the site-saturation mutagenesis. The primers were listed in Supporting Information Table S4. All mutants were confirmed by sequencing at Beijing Genomics Institute (BGI, China). The BD-EcaICE1_T3_ and its mutants were used for further Y2H and β-galactosidase activity assays.

### 2.7. Assay of β-galactosidase activity

The β-galactosidase activity was measured based on protocols from the yeast β-galactosidase assay kit manual (Thermo, USA). Single yeast colonies grown on SD/-LW medium of BD-EcaICE1_T3_ and its mutants with AD-EcaHOS1 were picked and transferred into 5 mL YPDA liquid medium, and incubated at 30 °C with 200 rpm shaker for 10-14 h, and the OD measurement at 660 nm was recorded. 1.0 mL of the culture medium was centrifuged at 13,000 g for 1 min, and the supernatant was removed. Then, 250 μL Y-PER and 250 μL ONPG solution were added, and immediately incubated at 37° C. When the mixed solution turned yellow, 200 μL stop solution was added to stop the reaction, and recorded the reaction time. After centrifugation at 13,000 g for 30 s, 200 μL supernatant was measured for the OD measurement at 420 nm. Each assay replicated three technological repeats in separate experiments. β-galactosidase activity was calculated as follows:

> β-Galactosidase (units)=(1000×OD_420_)(T×V×OD_660_)

where OD_420_, T, V, and OD_660_ were the OD measurement at 420 nm, reaction time (min), reaction solution volume (mL), and the OD measurement at 660 nm, respectively.

### 2.8. Data statistical analysis

The data values presented were the means ± standard errors (SE) of three replicates through statistical analysis via R software (v 3.5.1), and further analyzed by ANOVA and Duncan’s multiple range test to compare the differences between treatments at the *P* < 0.05 level.

## 3. RESULTS

### 3.1. Sequence Analysis of EcaICE1 and EcaHOS1

The sequence of *EcaICE1* gene was identical to our precious result [16]. The typical conserved domains, such as MYC-like bHLH domain, zipper structure at the C-terminus, S-rich (Serine-rich) acidic domain, SUMO binding site and NLS (Nuclear localization sequence) box, were found in EcaICE1 protein from the multiple alignments (Figure S1). Nevertheless, only *Eucalyptus* ICE1 proteins had an additional Q-rich (Glutamine-rich) domain, suggesting that the characteristics of ICE1 proteins might be different between *Eucalyptus* and the other plants.

The sequence of *EcaHOS1* gene was 2889 bp long and encoded a complete coding frame consisting of 962 amino acids. BLAST analysis revealed that EcaHOS1 shared a high sequence identity with other plant HOS1-like proteins, such as *A. thaliana* (52%, OAP11605), *Vitis viinifera* (60%, NP_001268014) and *Poncirus trifoliate* (58%, XP_024445010). The multiple alignments of plant HOS1 protein sequences (Figure 1) showed that the EcaHOS1 protein had a conserved RING finger domain and ELYS domain, similar to other plant HOS1 proteins. The conserved RING finger domain is the crucial functional region of E3 ubiquitin ligases [27]. These results showed that EcaHOS1 was the HOS1 protein and a new E3 ubiquitin ligase from *E. camaldulensis*, which may have a functional role in the ubiquitination pathway.

**Figure 1.**
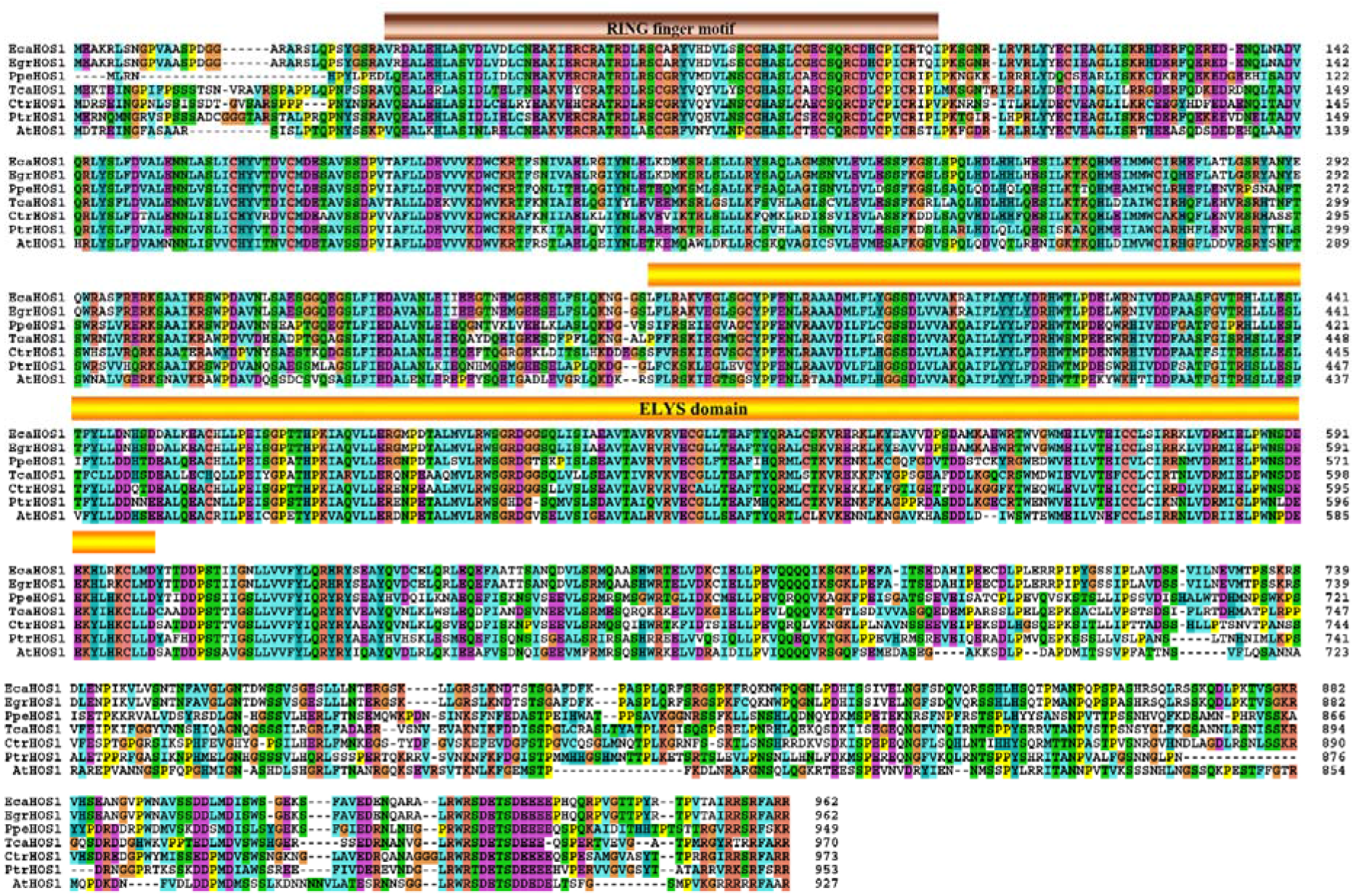
Amino acid alignment of the EcaHOS1 protein with six plant HOS1 proteins. The predicted protein domains were shown. The *Arabidopsis thaliana* AtHOS1 (Accession: AAP14668), *E. camaldulensis* EcaHOS1 (Accession: MH899181), *E. grandis* EgrHOS1 (Accession: XP_010055472), *Citrus trifoliata* CtrHOS1 (Accession: ACY92092), *Populus trichocarpa* PtrHOS1 (Accession: XP_024445010), *Prunus persica* PpeHOS1 (Accession: XP_020419310), *Theobroma cacao* TcaHOS1 (Accession: EOY24269) and *Vitis vinifera* VvHOS1 (Accession: NP_001268014) proteins are included.

### 3.2. Subcellular localization of EcaICE1

The multiple alignments (Figure S1) showed that there was an NLS box in the EcaICE1, implying that it may be nuclear localized protein. To confirm the result, the subcellular localization assay of EcaICE1 was carried out using a fusion protein of *EcaICE1* and *YFP* reporter gene, driven by 35S promoter. The yellow fluorescence of EcaICE1-YFP fusion protein was detected in the nucleus (Figure 2). These results indicated that EcaICE1 was a nuclear protein, similar to other woody plant ICE1 proteins [17,18,19].

**Figure 2.**
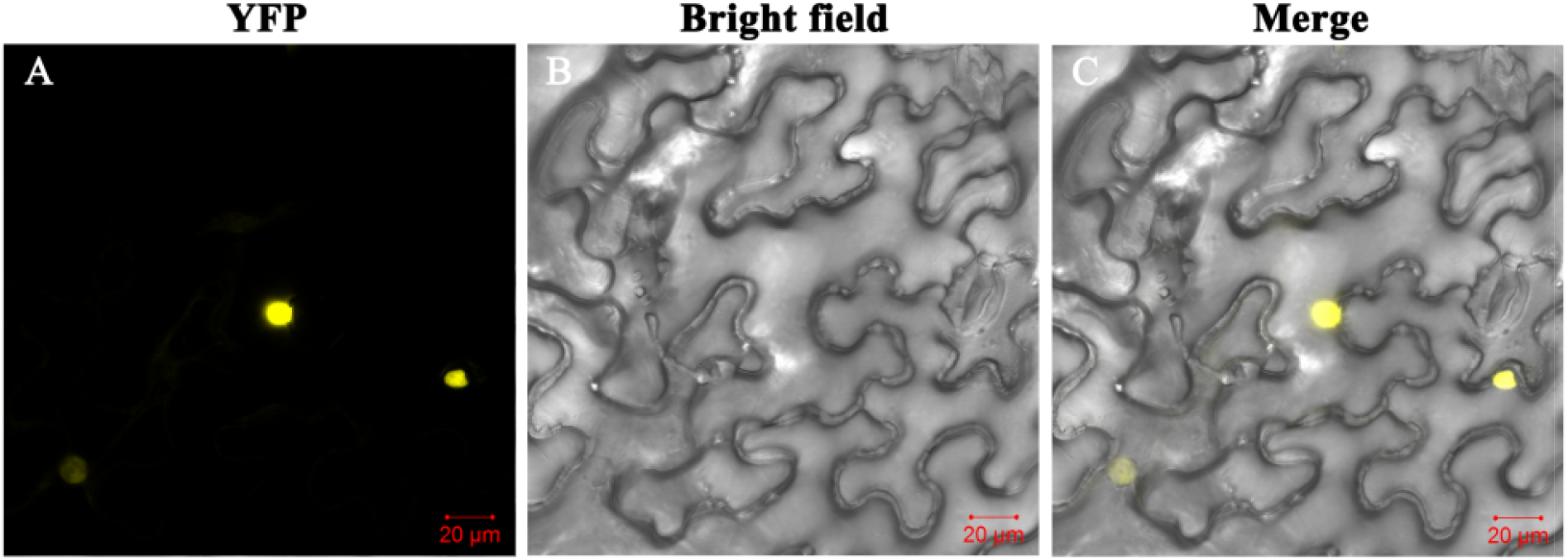
Subcellular localization of EcaICE1. A: the fluorescence signal plot of EcaICE1 in the dark field, B: the plot of EcaICE1 in the bright field, C: the plot of dark field and bright field superposition. Bar=20μm.

### 3.3. EcaICE1 could interact with EcaHOS1

Previous report showed that HOS1 could interact with ICE1 and mediate the ubiquitination of ICE1 both *in vitro* and *in vivo* in *A. thaliana*, which attenuated cold stress response by the ubiquitination/proteasome pathway [9]. To elucidate whether EcaHOS1 could also interact with EcaICE1 in *E. camaldulensis*, a BiFC assay was performed to identify the protein-protein interactions in tobacco leaves. Microscopic visualization results (Figure 3) revealed that there was no YFP fluorescent signal for the negative controls including EcaICE1-pSPYCE co-expressed with unfused pSPYNE or EcaHOS1-pSPYNE co-expressed with unfused pSPYCE whereas the YFP fluorescence was observed exclusively in the nucleus for the EcaICE1-pSPYCE co-transfected with EcaHOS1-pSPYNE. These results showed that EcaICE1 could interact with EcaHOS1 to form heterodimers at the nucleus.

**Figure 3.**
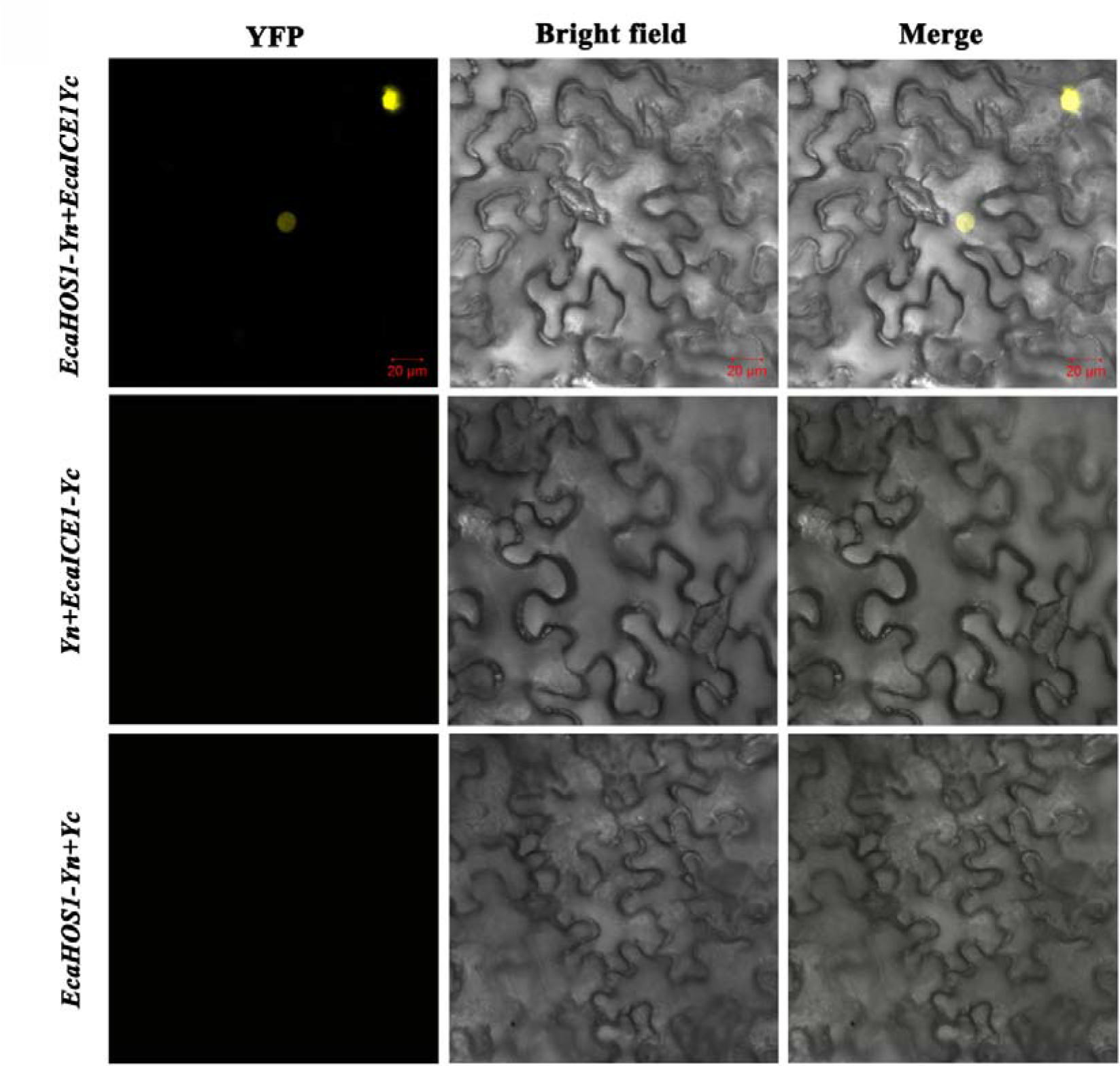
Bimolecular fluorescence complementation results of EcaICE1 and EcaHOS1. p35S::EcaHOS1-Yn:p35S::EcaICE1-Yc were co-expressed in tobacco cells; p35S::Yn:p35S::EcaICE1-Yc and p35S::EcaHOS1-Yn:p35S::Yc served as negative controls. The first column is the dark field fluorescence signal plot, the second column is the bright field plot, and the third column is the dark field and bright field plot. Bar=20μm.

### 3.4. Protein-protein interaction region between EcaICE1 with EcaHOS1

We perform the Y2H assay to further confirm the protein-protein interaction between EcaICE1 with EcaHOS1 and discover the key interaction region of EcaICE1 for the interaction. Unfortunately, both EcaICE1 and EcaHOS1 had autoactivation activity (Figure S3). Then, transcriptional activation assay demonstrated that amino acids from positions 84 to 125 in EcaICE1 were critical for the transactivation activity of EcaICE1 (Figure S2). Now the truncated EcaICE1 protein without transcriptional activation activity was constructed into vector pGBKT7, and co-transformed with AD-EcaHOS1 into yeast strain separately to find the protein-protein interaction region. The results showed that only the EcaICE1_T3_ interacted with EcaHOS1 and the other regions did not (Figure 4A), while all of these four truncated EcaICE1 proteins could interact with AtHOS1(Figure 4B), indicating that the N-terminus region of EcaICE1(126-185aa) was the key region for its interaction with EcaHOS1 protein, quite different from *A. thaliana* [28].

**Figure 4.**
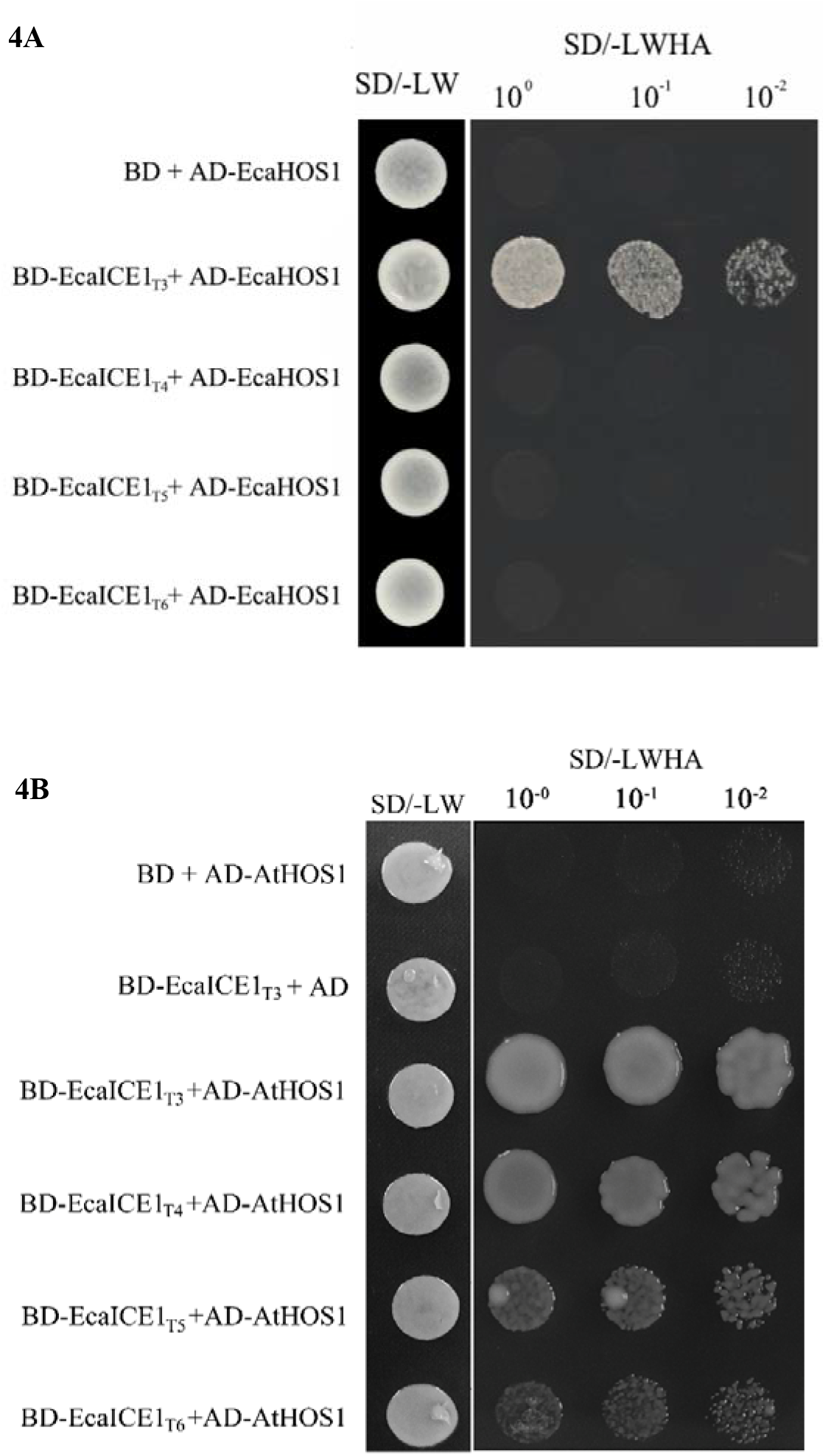
The interaction identification of truncated EcaICE1 with EcaHOS1 (4A) and AtHOS1 (4B) by yeast two-hybrid methods. BD: pGBKT7 vetor; AD: pGADT7 vetor; SD/-LW: SD/-Leu-Trp: double dropouts, SD medium with -Leu/-Trp; SD/-LWHA: quadruple dropouts, SD medium with-Leu-Trp-His-Ade; 10^0^, 10^-1^, 10^-2^: dilutions with 1, 10 and 100 times respectively.

### 3.5. Effects of key phosphorylation site of EcaICE1 for its interaction with EcaHOS1 protein

We searched the phosphorylation sites within the N-terminus region (126-185 aa) of EcaICE1 by bioinformatics software NetPhos 3.1 and found that Ser 143 (Ser at 143 aa), Thr 145(Thr at 145 aa), Ser 158(Ser at 158 aa) and Ser 184(Ser at 184 aa) were predicted as the potential phosphorylation sites. After the substitution of these four sites by Alanine (S143A, T145A, S158A and S184A, respectively) using site-direct mutagenesis based on the EcaICE1_T3_ (abbreviated as T3), Y2H results (Figure 5A) showed that only T3(S158A) blocked its interaction with EcaHOS1, while the residual three mutants and T3 still worked. It is interesting that both T3 and its mutants of EcaICE1 could interact with AtHOS1 (Figure 5B), indicating that the ubiquitination pathway of ICE1 by HOS1 may be different between *Eucalyptus* and *Arabidopsis*. β-galactosidase assay (Table 2) revealed that β-galactosidase activity of T3(S158A) was not significant from negative control at *P*<0.05. The β-galactosidase activity of T3(T145A) and T3(S184A) was not significant from wild type T3 at *P*<0.05, while that of T3(S143A) was significantly higher than wild type T3 (*P*<0.05). The β-galactosidase activity assay also confirmed that only T3(S158A) blocked the protein interactions between EcaICE1 and EcaHOS1. These results suggested that Ser 158 was the key phosphorylation site of EcaICE1 for its interaction with EcaHOS1.

**Figure 5.**
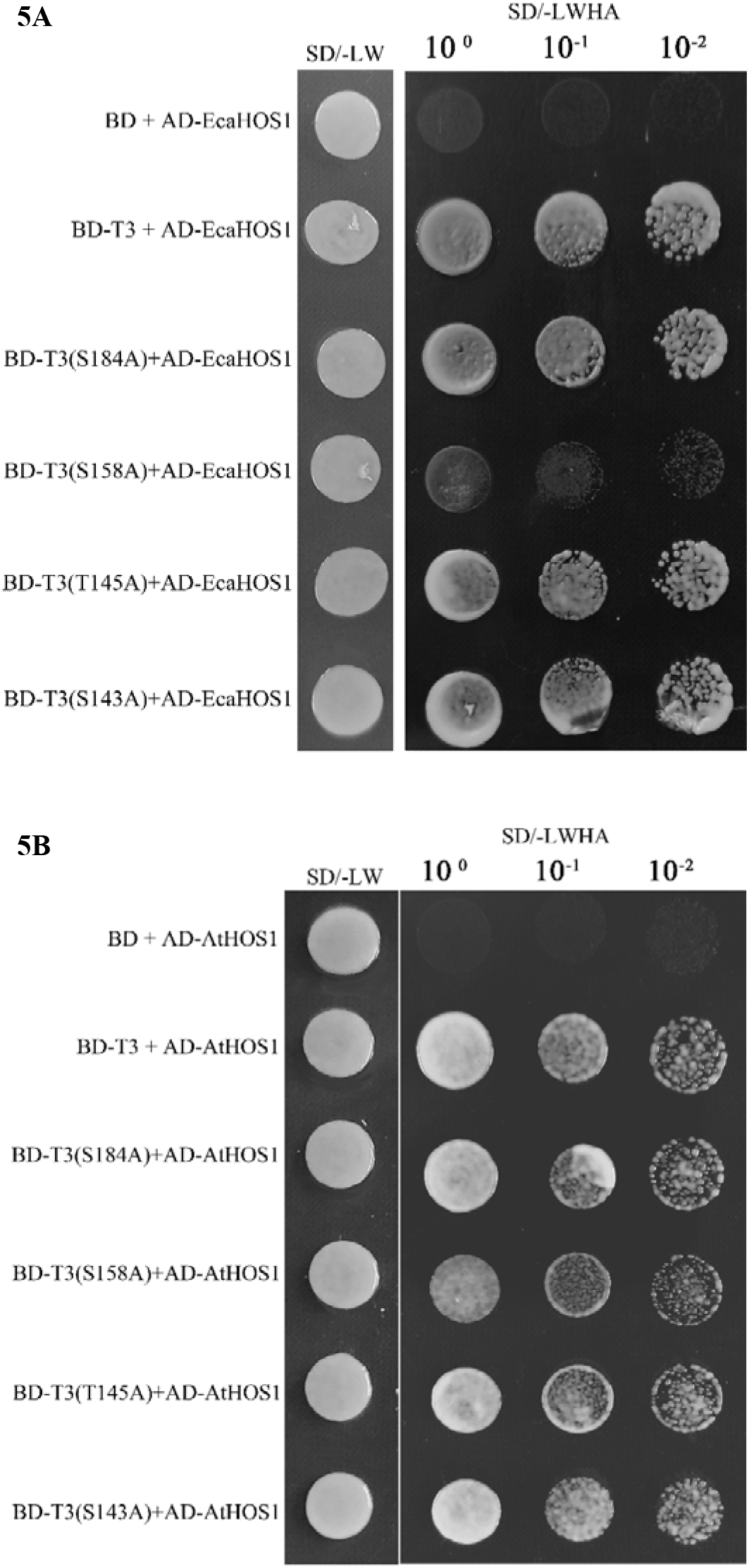
The interaction identification of EcaICE1-T3 and its mutants with EcaHOS1 (5A) and AtHOS1 (5B) by yeast two-hybrid methods. BD: pGBKT7 vetor; AD: pGADT7 vetor; T3: truncated EcaICE1-T3; T3(S143A), T3(T145A), T3(S158A), T3(S184A): the mutant of truncated EcaICE1-T3 at serine 143, threonine 154, serine 158 and serine 184 by alanine, respectively; SD/-LW: SD/-Leu-Trp: double dropouts, SD medium with -Leu/-Trp; SD/-LWHA: quadruple dropouts, SD medium with-Leu-Trp-His-Ade; 10^0^, 10^-1^, 10^-2^: dilutions with 1, 10 and 100 times respectively.

**Table 2.** Measurement of β-galactosidase activity of protein interactions of EcaICE1-T3 and its mutants with EcaHOS1.

| BD vector | AD vector | Yeast cells<br>on SD/-LW | Yeast cells<br>on SD/-LWHA | β-galactosidase activity |
| --- | --- | --- | --- | --- |
| T3(S143A) | EcaHOS1 | White colonies | White colonies | 27.29±2.72 a |
| T3(T145A) | EcaHOS1 | White colonies | White colonies | 13.29±0.31 b |
| T3(S158A) | EcaHOS1 | White colonies | None | 5.90±1.74 c |
| T3(S184A) | EcaHOS1 | White colonies | White colonies | 13.56±0.15 b |
| T3 | EcaHOS1 | White colonies | White colonies | 12.32±0.16 b |
| BD | AD | White colonies | None | 5.16±1.35 c |

Note: Values for β-galactosidase activity followed by the same letter are not significantly different at $\alpha = 0.05$ using Duncan's test.
BD: pGBKT7 vector; AD: pGADT7 vector; T3: truncated EcaICE1-T3; T3(S143A), T3(T145A), T3(S158A), T3(S184A): the mutant of truncated EcaICE1-T3 at serine 143, threonine 154, serine 158 and serine 184 by alanine, respectively; SD/-LW: SD/-Leu-Trp: double dropouts, SD medium with -Leu/-Trp; SD/-LWHA: quadruple dropouts, SD medium with -Leu-Trp-His-Ade.

## 4. Discussion

*Eucalyptus* can increase freezing tolerance by cold acclimation as well as other plant species. *CBF* genes have been cloned from *Eucalyptus* [20,21,22,25] and their overexpression in cold-sensitive *Eucalyptus* could improve the freezing tolerance [23]. Cao et al. further reported that there were 17 CBF orthologs in the *E*. *grandis* genome and that 14 CBF genes were located on the scaffold 1 within a cluster of about 117 Kb [24]. We first cloned an ICE1 gene *EcaICE1* from *E. camaldulensis* and found that ectopic expression of *EcaICE1* could confer improved cold tolerance and increase the expression level of downstream genes in transgenic tobaccos [16]. However, the factors controlling ICE1 protein stability associated with cold stress response in *Eucalyptus*, that is, whether there is similar ubiquitination or SUMOylation pathway of ICE1 to *Arabidopsis*, are still lack of empirical studies. In this current study, we further researched the ubiquitination pathway of ICE1 mediated by HOS1 in *Eucalyptus* and characterized EcaICE1 protein having a direct physical interaction with EcaHOS1 *in vivo* (Figure 3, 4A, and 5A).

E3-ubiquitin-ligases-mediated ubiquitination are emerging as major regulators in plants’ response to abiotic stress, circadian rhythm control, cell cycling and plant-microbe interactions [29,30,31]. High expression of osmotically responsive genes1 (HOS1), one of the RING finger-containing E3 ubiquitin ligase, was initially described as a cold-signaling attenuator of ICE1 [9,32], and also involved in ethylene signal transduction [33] and photoperiodic flowering [34] in *Arabidopsis*. It is reported that most of the plant species just have a single copy of the *HOS1* gene [35], suggesting that HOS1 would have important roles in various plant species. Based on the multiple alignments of EcaHOS1 with other plant HOS1 proteins (Figure 1), the RING finger domain is highly conserved among the presented plants, and the first residue in the RING finger of all seven HOS1 proteins is Leu, which is Cys in animal inhibitor of apoptosis (IAPs) [36]. The RING-finger domain is the crucial player in the ubiquitin-dependent protein degradation system [37], indicating that plant HOS1 proteins are highly conserved E3 ubiquitin ligase. In addition, all seven HOS1 proteins have an highly conserved embryonic large molecule derived from yolk sac (ELYS) domain, which is required for the protein recruitment of the nuclear pore complex (NPC), implying that HOS1 also has nonproteolytic roles such as mRNA export and chromatin remodeling [35]. These analyses indicate that *EcaHOS1* is the HOS1 gene from *E. camaldulensis*, and its encoded protein may have E3 ubiquitin ligase activity and mediate ubiquitination in the nucleus.

BiFC assay showed that EcaICE1 could interact with EcaHOS1 in nucleus, similar to *A. thaliana* [9], but which region of EcaICE1 was the key region for the protein-protein interactions is still unknown. Therefore, using different truncated EcaICE1 without autoactivation activity as bait protein, the N-terminal 126-185 amino acid of EcaICE1 protein was further identified as the key region by Y2H assay. Interestingly, the key region of EcaICE1 (126-185 aa) is not located within the conserved bHLH domain, the zipper motif or ACT-like domain at the C-terminus region of EcaICE1. However, in *Arabidopsis*, AtHOS1 interacted with the C-terminus of AtICE2 containing the zipper motif [28]. Surprisingly, all truncated EcaICE1 could interact with AtHOS1, indicating that the zipper motif at C-terminus of ICE protein is necessary for its interaction with AtHOS1.

A number of evidences show that non-lysine residues, such as serine, threonine and cysteine, are ubiquitylation sites of many E3 ubiquitin ligases [38,39,40]. Miura et al. reported that substitution of Ser 403 by Alanine (Ala) in *Arabidopsis* ICE1 could inhibit the polyubiquitylation of ICE1 *in vivo*, but did not affect the degradation of ICE1 [41]. They argued that although Ser 403 is not the main target residue for ubiquitylation or SUMOylation, it is a key residue for the attenuation of cold-stress responses by HOS1-mediated degradation of ICE1 [41]. Here, we further found that Ser 143, The 145, Ser 158 and Ser 184 within 126-185 aa of EcaICE1 were predicted as the potential phosphorylation sites based on bioinformatics method. Only substitution of Ser 158 by Ala (S158A) using site-direct mutagenesis blocked the protein-protein interactions between EcaICE1 and EcaHOS1 using Y2H assay (Figure 5A), and β-galactosidase activity assay (Table 2) also confirmed this result. Surprisingly, all of EcaICE1 mutants could interact with AtHOS1 again (Figure 5B), indicating that the conserved domains at the C-terminus of ICE indeed are necessary for its interactions with AtHOS1 and that the ubiquitylation site of ICE for AtHOS1 maybe the Ser 403. These results suggested that ubiquitination pathway of ICE1 mediated by HOS1 may be different between *Eucalyptus* and *Arabidopsis* and that Ser 158 was the key phosphorylation site of EcaICE1 for its interaction with EcaHOS1. As for whether EcaHOS1 acts as an E3 ubiquitin protein ligase and whether it could mediate the degradation of EcaICE1 for playing a role in cold stress response in *Eucalyptus*, including whether Ser 158 affects the EcaHOS1-mediated ubiquitination of ICE1, need further experiments to investigate.

## 5. Conclusions

Herein, we amplified the gene *EcaICE1* and *EcaHOS1* from *E. camaldulensis*, and the *EcaICE1* sequence was the same as the previous result, and the deduced EcaHOS1 protein was highly conserved with other plant HOS1 proteins. EcaICE1 was located in nucleus, and it could interact with EcaHOS1 in nucleus by BiFC assay. Moreover, Y2H assays confirmed the interaction between EcaICE1with EcaHOS1 and revealed that the N-terminal region from position 126 to 185 in EcaICE1 was the key region for its interaction with EcaHOS1 protein. Finally, Ser 158 was the key phosphorylation site of EcaICE1 for its interaction with EcaHOS1 by Y2H and β-galactosidase assay using site-direct mutagenesis. Taken together, we made a foundation on the ubiquitination regulation mechanism of EcaICE1 mediated by EcaHOS1.

## Supplementary Materials

The following are available online at www.mdpi.com/, Table S1: Primers used in the subcellular localization experiments; Table S2: Primers used in the BiFC experiments; Table S3: Primers used in the Y2H experiments; Table S4: Primers used in the site-direct mutagenesis; Figure S1: Amino acid alignment of the EcaICE1 protein with other plant ICE1 proteins; Figure S2: Transcriptional activation region analysis of EcaICE1 protein; Figure S3: Transcriptional activation analysis of EcaICE1 and EcaHOS1protein.

## Author Contributions

L.Y. conceived and designed the experiments; C.L., Z.W. and Z.Z. performed the experiments; Z.Z. and L.Y. analyzed the data; C.L. and L.Y. wrote the paper. C.L., Z.W. and H.J. contributed equally to this research.

